# JIP4 is recruited by the phosphoinositide-binding protein Phafin2 to promote recycling tubules on macropinosomes

**DOI:** 10.1101/2020.09.29.319111

**Authors:** Kia Wee Tan, Viola Nähse, Coen Campsteijn, Andreas Brech, Kay Oliver Schink, Harald Stenmark

## Abstract

Macropinocytosis allows cells to take up extracellular material in a non-selective manner. The molecular mechanisms that mediate recycling of membranes and transmembrane proteins from macropinosomes still need to be defined. Here we report that JIP4, a coiled-coil containing protein previously described to bind to microtubule motors, is recruited to retromer- and actin-containing tubulating subdomains on macropinosomes by binding to the PH domain of the phosphatidylinositol 3-phosphate (PtdIns3P)-binding protein Phafin2. This recruitment is not shared by the closely related isoforms JIP3 and Phafin1. Disruption of Phafin2 or PtdIns3P impairs JIP4 recruitment to macropinosomes whereas forced localization of Phafin2 to mitochondria causes mitochondrial targeting of JIP4. While knockout of JIP4 suppresses tubulation, overexpression enhances tubulation from macropinosomes. JIP4 knockout cells display increased retention of macropinocytic cargo in both early and late macropinosomes, consistent with a recycling defect. Collectively, these data identify JIP4 and Phafin2 as components of a tubular recycling pathway that operates from macropinosomes.

## Introduction

Macropinocytosis is a process that enables cells to take up large amounts of extracellular fluid [1]. This fluid is internalized into large vesicles which are called macropinosomes. During this process, large regions of plasma membrane and the proteins within are internalized. In order to preserve the composition of the plasma membrane, it is important that membranes and plasma membrane proteins are recycled and transported back to the cell surface.

Directly after internalization, macropinosomes frequently tubulate and bud off small vesicles [2]. This process, sometimes called “piranhalysis”, has frequently been observed in cells [3, 4], but the underlying mechanisms are poorly understood. Tubulation from vesicle membranes often requires the action of membrane-bending proteins such as sorting nexins [5]. One of these sorting nexins, SNX5, has been shown to regulate macropinocytosis [6, 7]. In addition, tubulation and the formation of vesicles typically require motor proteins which exert pulling forces on the nascent membrane tubule. Often, multiple motor proteins are involved in a “tug of war”, and by this generate forces which drive scission of the membrane [8].

This motor-driven tubule pulling and scission requires adaptor proteins, which link motor proteins to the tubule membrane. JIP4 is a coiled-coil protein which can bind to both dynein and kinesin motor proteins [9, 10]. It can also bind to the small GTPase ARF6 [11]. This binding has been proposed to control a motor switch which controls endocytic recycling during cytokinesis [9]. ARF6 and JIP3/JIP4 have also been shown to regulate endosomal recycling of the matrix metalloproteinase MT1-MMP [12]. The transmembrane protein TMEM55B recruits JIP4 to lysosomes to mediate long-distance lysosome transport [13]. This is especially important in neurons, and mutations in the *Drosophila melanogaster* homolog *sunday driver* affect axonal long distance transport [14]. Moreover, a recent preprint showed that tubulating lysosomes contain JIP4 [15].

Here, we show a novel role of JIP4 on tubulating macropinosomes. We show that the lipid-binding protein Phafin2 recruits JIP4 to retromer-containing tubules of tubulating macropinosomes in a phosphatidylinositol 3-phosphate (PtdIns3P)-dependent fashion. Deletion of JIP4 reduces tubulation from macropinosomes, accompanied by retention of fluid-phase cargo in early and late macropinosomes. Conversely, overexpression of both JIP4 and its recruiter Phafin2 leads to strongly enhanced tubulation. These results suggest that JIP4 is important for membrane recycling from newly-internalized macropinosomes by promoting membrane tubulation.

## Results

We have recently identified the phosphoinositide-binding protein Phafin2 as a novel regulator of macropinosome formation [16]. Using a two-hybrid screen for Phafin2 interactors, we identified the protein JIP4 as a potential interactor of Phafin2 (Supplementary Table S1). This was interesting since JIP4 and its homolog JIP3 have been implicated in macropinocytosis [17], although their function has remained largely unknown.

We first confirmed the interaction of JIP4 with Phafin2 using yeast two-hybrid interaction assays with truncation mutants of Phafin2 against the identified interaction region in JIP4. Phafin2 contains a PH and a FYVE domain, which are both involved in lipid binding (Figure 1A). JIP4 interacts with Phafin2 only via the Phafin2 PH domain (Figure 1B), as deletion of the PH domain, but not the FYVE domain abolished expression of the reporter gene. To extend these results to mammalian cells, we performed proximity biotinylation labeling using cell lines stably expressing APEX2-fusions of full length or deletion mutants of Phafin2, with cell lines expressing cytosolic or membrane anchored APEX2 serving as negative controls. Semi-quantitative mass spectrometry analysis showed that deletion of the PH domain of Phafin2 greatly impaired biotinylation of JIP4, while deletion of the FYVE domain, which is required for localization of Phafin2 to early macropinosomes, [16] did not (Figure 1C, Supplementary Table S2). Together, these experiments indicate that the FYVE domain of Phafin2 is not involved in the interaction with JIP4 and that a local membrane environment is not required.

**Figure 1:**
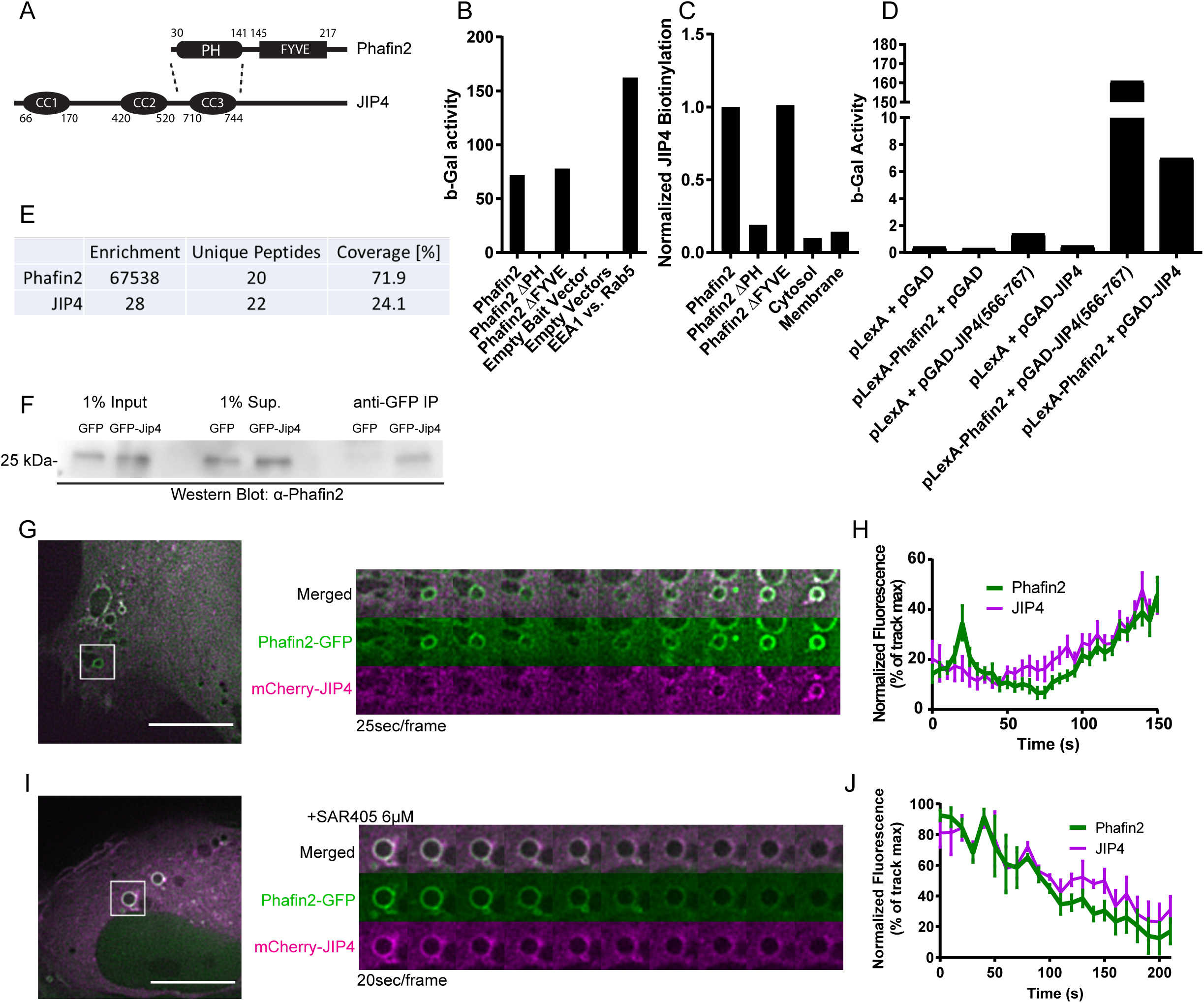
JIP4 interacts and colocalizes with Phafin2. A) Domain structure of JIP4 and Phafin2, dotted lines indicate interacting regions. CC1, 2 and 3 indicate predicted coiled coils. B) β-galactosidase activity derived from yeast two-hybrid assay expressing the specified constructs, with JIP4 (566-767aa) as prey. C) Biotinylated JIP4 detected in mass spectrometry following labeling with the specified APEX2 fusion constructs, normalized to wildtype Phafin2. Cytosol is a control consisting only of the soluble APEX2, while Membrane is a control consisting of the APEX2 fused to a signal peptide that targets it primarily to the plasma membrane. D) β-galactosidase activity derived from yeast 2-hybrid assay expressing the specified constructs, with full length Phafin2 as bait. E) Endogenous JIP4 detected in mass spectrometry following affinity purification of tagged Phafin2, fold change over control cells expressing only the affinity tag. F) Immunoprecipitation of GFP-JIP4 with GFP-Trap, western blotting against endogenous Phafin2 in RPE1 lysate. Uncropped blot in Suppl. Fig 1. Representative of 3 independent experiments. G) RPE1 cell expressing Phafin2-GFP and mCherry-JIP4, imaged live. Montage gallery of boxed region. H) Mean fluorescence measurements along the limiting membrane of macropinosomes, each measurement normalized to the mean of the individual time series, aligned at timepoint 15sec to the burst of Phafin2 fluorescence on nascent macropinosomes, +/-SEM (n=13 macropinosomes). I) RPE1 cell expressing Phafin2-GFP and mCherry-JIP4, treated with SAR405 (VPS34 inhibitor) to remove PtdIns3P from macropinosomes, imaged live. Montage gallery of boxed region. J) Mean fluorescence measurements along the limiting membrane of macropinosomes treated as in I, each measurement normalized to the mean of the individual time series +/-SEM (n=4 macropinosomes). Scale bars in (G) and (I) are 10µm.

To verify that full length JIP4 was also capable of interacting with Phafin2, we used yeast two-hybrid assays and immunoprecipitation. Full length JIP4, like the isolated interaction region previously identified, triggered expression of the reporter gene in the yeast two-hybrid assay (Figure 1D). To assess the interaction between Phafin2 and JIP4 in mammalian cells, we performed tandem affinity purification using lysates from RPE1 cells stably expressing Localization and Affinity Purification (LAP) tagged Phafin2. Semi-quantitative mass spectrometry analysis identified JIP4 as a strong interactor in these pulldowns, with a 28-fold enrichment for JIP4 compared to control cells expressing solely the LAPtag (Figure 1E, Supplementary Table S3). Conversely, we precipitated GFP-JIP4 from cell lysate of RPE1 cells stably expressing GFP-JIP4 using GFP-TRAP magnetic beads. Immuno-blotting with an anti-Phafin2 antibody showed that endogenous Phafin2 was co-precipitated with GFP-JIP4, but not with GFP alone (Figure 1F, Suppl. Fig. 1D).

We used live-cell microscopy to assess if Phafin2 and JIP4 localize to similar cellular structures. Phafin2 shows a biphasic localization to macropinosomes, one to nascent macropinosomes directly after scission from the membrane and one to macropinosomes that have matured into endosome-like vesicles (in this study we will refer to these as early macropinosomes, as they acquire markers of early endosomes) [16]. Interestingly, we found that JIP4 selectively co-localizes with Phafin2 at early macropinosomes but did not co-localize with Phafin2 on nascent macropinosomes (Figure 1G, H, Supplementary Video 1). This could suggest that a binding site of JIP4 is not accessible on newly-formed vesicles. Phafin2 requires PtdIns3P, generated by the PtdIns 3-kinase VPS34, to localize to early macropinosomes [16]. To test if macropinosome localization of JIP4 is dependent on Phafin2, we treated cells with the selective VPS34-inhibitor SAR405 [18] and assessed JIP4 localization. Addition of SAR405 led to a rapid displacement of both Phafin2 and JIP4 from the membrane (Figure 1I, J), suggesting that JIP4 depends on Phafin2 for the macropinosome localization.

As a putative recruiter, modulation of Phafin2 protein levels by overexpression or ablation would be expected to affect JIP4 localization. We assessed endogenous JIP4 localization to early endosomes in wild-type, Phafin2 KO [16] or Phafin2 overexpressing RPE1 cells by immunostaining for JIP4 and the early-endosomal antigen EEA1 and quantifying JIP4 intensity in EEA1-labelled endosomes. We found that JIP4 showed reduced localization to early endosomes if Phafin2 was deleted. In contrast, overexpression of Phafin2 led to a strong recruitment of JIP4 to EEA1-positive endosomes (Figure 2A, B).

**Figure 2:**
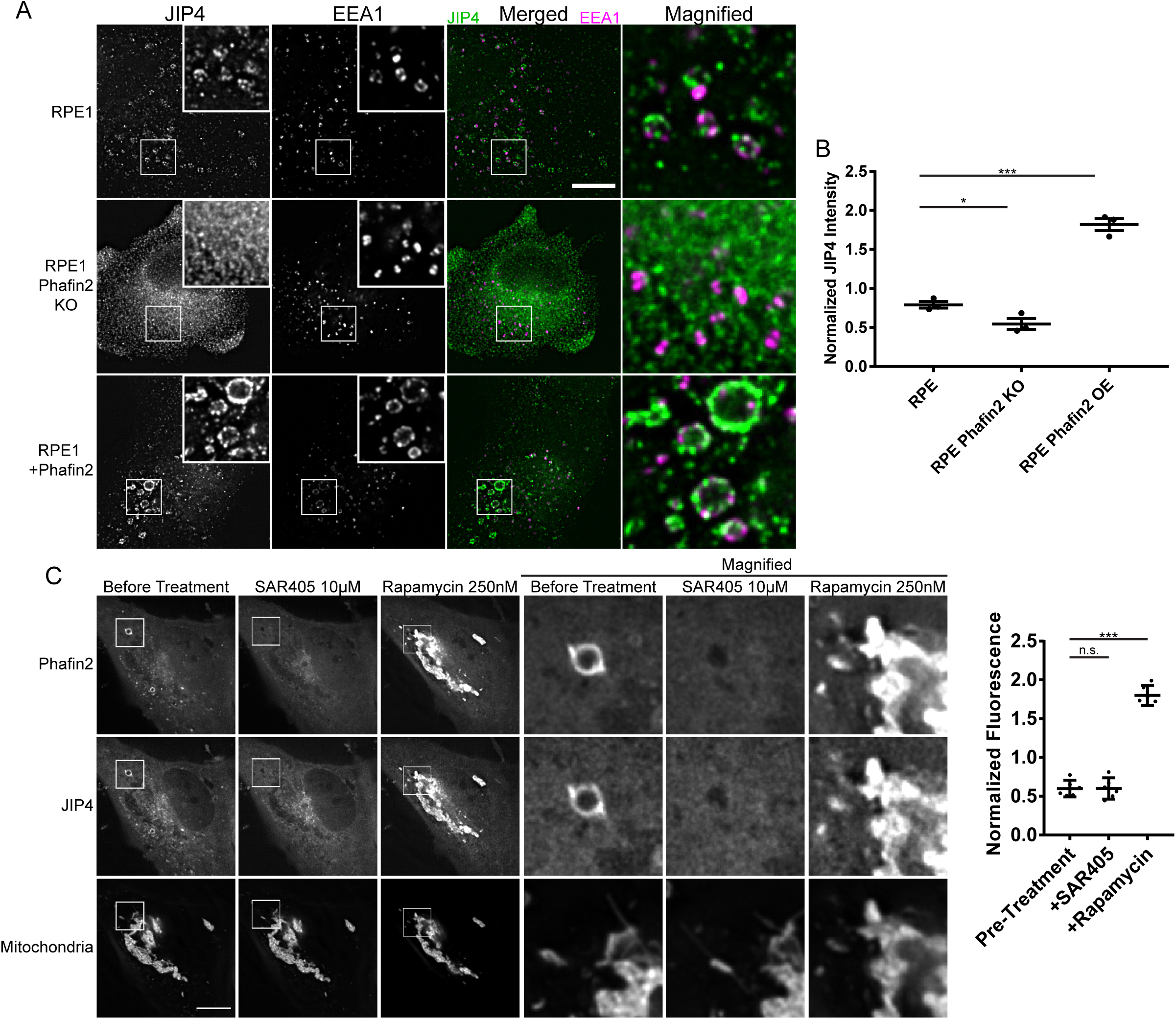
Membrane recruitment of JIP4 by Phafin2. A) Representative images of RPE1 cells of the specified genotypes, fixed and immunostained against JIP4 and EEA1. Brightness settings are equal across all images and magnifications. Scale bar is 10 µm. B) Mean intensities of JIP4 immunostaining inside EEA1 positive vesicles, each experiment normalized against mean of all datapoints in that experiment. Mean of 3 experiments shown, +/- s.e.m. (3530-6121 vesicles per condition per experiment) C) RPE1 cell expressing Phafin2-FRB-mNeonGreen, mCherry-JIP4, and a mitochondrial-anchored 2xFKBP. Shown are images of the same cell before addition of 10µM SAR405, after SAR405 has removed macropinosome PtdIns3P, and after 250nM rapamycin has recruited Phafin2 to the mitochondrial membrane. Scale bar is 10µm. D) JIP4 fluorescence at the mitochondria, images acquired of the same cells under the three sequential conditions, segmented and measured using the mitochondrial marker as shown in C). Error bars are 95% C.I. (n=6 cells)

To further support that JIP4 is recruited by Phafin2, we used a chemical dimerization system to redirect Phafin2 to mitochondria and monitored the localization of JIP4. To this end, we expressed an FRB and fluorophore tagged Phafin2, a mitochondrially anchored 2xFKBP domain (Tom70-mTagBFP2-2xFKBP), and a fluorophore tagged JIP4 in RPE1 cells. FKBP and FRB domains heterodimerize in the presence of rapamycin [19], allowing redirection of FRB-tagged Phafin2 to the mitochondria by adding rapamycin to the extracellular solution. Cells expressing all three components were first treated with SAR405, leading to a dissociation of Phafin2 and JIP4 from macropinosomes (Figure 2C, D). Addition of rapamycin caused FRB-tagged Phafin2 to be recruited to the mitochondria (Figure 2C, D). JIP4 was co-recruited with Phafin2 to the mitochondria, indicating that Phafin2 does not require additional co-factors to recruit JIP4. Taken together, these data indicate that interaction of JIP4 with Phafin2 is sufficient for its subcellular targeting.

Both Phafin2 and JIP4 have homologs in the human genome, Phafin1 and JIP3, which share a large degree of sequence homology (Figure 3A, B). It is often implied that JIP3 and JIP4 have similar functions [12, 17, 20, 21]. We therefore asked if they could functionally replace each other. First, we tested if Phafin1 or Phafin2 can bind to JIP3 using direct two-hybrid interaction assays. To this end, we isolated the region corresponding to the identified JIP4-Phafin2 interaction domain from JIP3 based on the JIP3/JIP4 sequence homology. We did not observe any interaction of either Phafin1 or Phafin2 with JIP3 (Figure 3C). We also tested if Phafin1 can bind to JIP4 by two-hybrid interaction assays. Despite the high sequence homology between the PH domains of Phafin1 and Phafin2 (Figure 3B), we did not observe any interaction between Phafin1 and JIP4 (Figure 3D). This suggests that the interaction between Phafin2 and JIP4 is specific.

**Figure 3:**
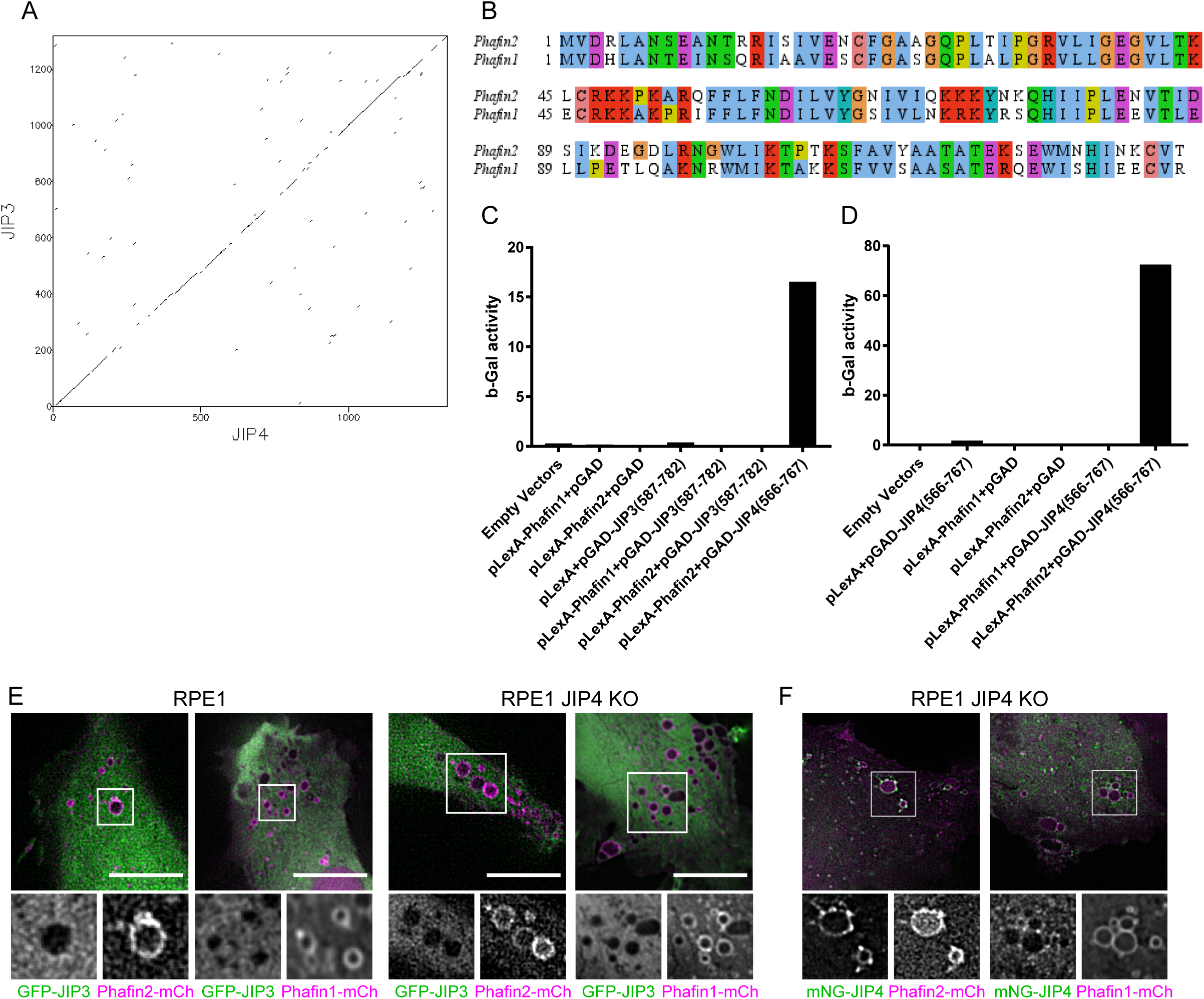
JIP4 and Phafin2 interaction is not shared with their respective isoforms. A) Dotplot of human JIP3 against human JIP4. Similarity matching using BLOSUM62 with a sliding window of 5 residues and a threshold score of 20 [33]. Note the unevenly distributed regions of high sequence conservation. B) Sequence alignment of the Phafin2 and Phafin1 PH domain. C) β-galactosidase activity derived from yeast 2-hybrid assay expressing the specified constructs. Representative of 3 independent experiments. D) β-galactosidase activity derived from yeast 2-hybrid assay expressing the specified constructs. Representative of 3 independent experiments. E) Representative images of cells of the indicated genotypes expressing mNeonGreen JIP3 and a Phafin isoform. JIP3 is not recruited to macropinosomes. Scale bars are 10µm. F) Representative JIP4 KO cell expressing mNeonGreen-JIP4 and a Phafin isoform. Phafin1 does not recruit JIP4.

We generated an RPE1 cell line deleted for JIP4 by CRISPR/Cas9 to facilitate further investigation. This cell line was verified by Sanger sequencing (Suppl. Fig 1A), immunoblotting (Suppl. Fig 1B) and immunostaining (Suppl. Fig 1C) and was used for all subsequent assays where a JIP4 KO is indicated.

To confirm the data obtained through two-hybrid interaction assays and to verify that the full length proteins do not contain interaction sites outside the two-hybrid tested regions, we coexpressed different combinations of Phafin1/2 and JIP3/4 in RPE1 cells. JIP3 and JIP4 dimerize through coiled-coil regions [10, 11, 22] and could form heterodimers in cells, and by this be recruited together. To account for this, we expressed GFP-tagged JIP3 together with either Phafin2 or Phafin1 in both wild-type cells and cells deleted for endogenous JIP4 and assayed JIP3 localization (Figure 3E). While Phafin1 – similarly to Phafin2 – localizes to macropinosomes, we did not observe any localization of JIP3 to these vesicles (Figure 3E). Conversely, we co-expressed mNeonGreen-JIP4 together with either Phafin2 or Phafin1 in cells deleted for endogenous JIP4. JIP4 was readily recruited to macropinosomes by Phafin2, but not by Phafin1 (Figure 3F). Taken together, these data show that Phafin2 interacts with JIP4. The Phafin2 homolog Phafin1 does not bind to JIP4, and the JIP4 homolog JIP3 is unable to bind to Phafin2.

As JIP4 localized specifically to early macropinosomes but not nascent macropinosomes, we next analysed JIP4 localization in relation to known early endosomal markers. To minimize the risk of overexpression artefacts, we generated a stable cell line expressing mNeonGreen-tagged JIP4 under control of the weak PGK promoter. We found that JIP4 localizes to early macropinosomes, labelled by the small-GTPase RAB5. However, JIP4 did not localize to the whole macropinosome, but was restricted to small, tubular subregions of the macropinosome (Figure 4A, B, Supplementary Video 2). In order to further characterize these structures, we expressed a marker of PtdIns3P-containing membrane tubules, a tandem FYVE domain of the protein WDFY2 [23], together with JIP4. JIP4 localized to mCherry-2xFYVE^(WDFY2)^ labelled tubules (Figure 4C, D). In contrast, in cells deleted for the JIP4 recruiter Phafin2, this localization was largely lost (Figure 4C, D). We then asked if Phafin2 shows a similar localization to tubular structures. Halo-tagged Phafin2 was expressed in cells at a very low level together with mNeonGreen-JIP4. Using these weakly expressing cells, we observed that Phafin2 labelled the limiting membrane of macropinosomes, but was enriched on tubular structures (Figure 4E). JIP4 showed only minimal staining of the limiting membrane and was strongly concentrated at Phafin2-labelled macropinosome tubules (Figure 4E).

**Figure 4:**
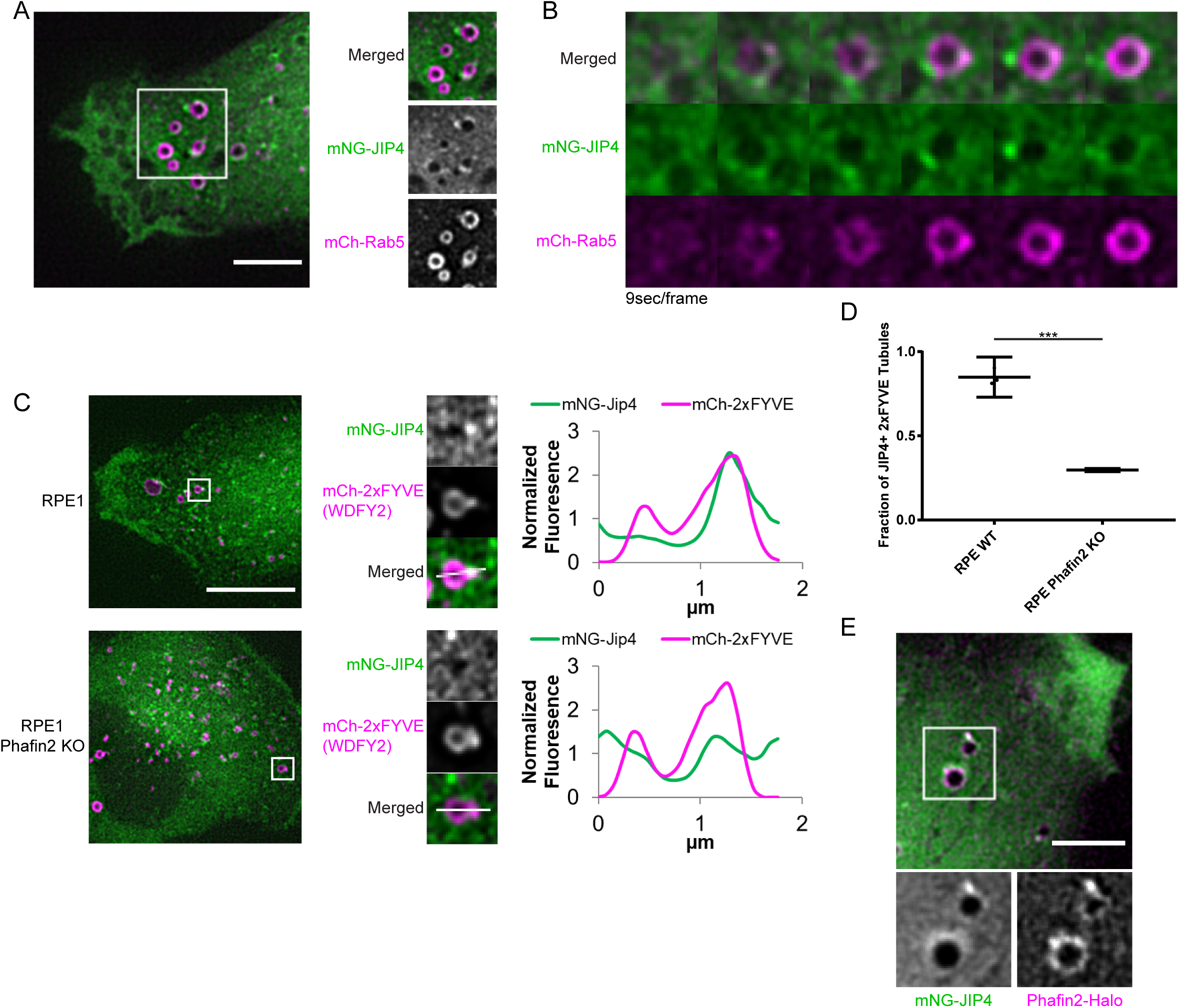
JIP4 is recruited to macropinosome tubules by Phafin2. A) Representative image of RPE1 cell expressing mNeonGreen-JIP4 and mCherry-Rab5, imaged live. B) Montage gallery of a macropinosome as it matures into a Rab5 positive early macropinosome and acquires JIP4. C) Representative images and magnifications of RPE1 cells of the specified genotype expressing mNeonGreen-Jip4 and mCherry-2xFYVE(WDFY2). Line plots are taken along the indicated line from left to right. Scale bar is 10 µm. D) Fraction of 2xFYVE(WDFY2) tubules per cell positive for mNG-JIP4. Positive threshold set at 1.5x cytoplasmic fluorescence. Mean of 3 experiments +/- s.e.m. shown. (42-105 events per condition per experiment) E) Representative image of an RPE1 cell expressing mNeonGreen-JIP4 and weakly expressing Phafin2-Halotag. Note that both Phafin2 and JIP4 stand out strongly against the diffuse cytoplasmic fluorescence on tubules. Scale bar is 5µm.

While these Phafin2 and JIP4 decorated structures resembled membrane tubules extruded from the limiting membrane of macropinosomes, the resolution of light microscopy cannot distinguish between organelle contact sites and emanating tubules. To verify that the JIP4-labelled tubules are continuous with the macropinosome membrane, we performed correlative light and electron microscopy (CLEM) using mNeonGreen-tagged JIP4. We first followed JIP4 localization together with Halo-2xFYVE^(WDFY2)^ by live cell imaging and then chemically fixed the cell during imaging (Figure 5A, B). Fixed cells were processed for electron microscopy and micrographs for electron tomography were collected (Figure 5C). Reconstruction of these tomograms showed that the JIP4-labelled tubules formed continuous structures with the limiting membrane of the macropinosome (Figure 5D).

**Figure 5:**
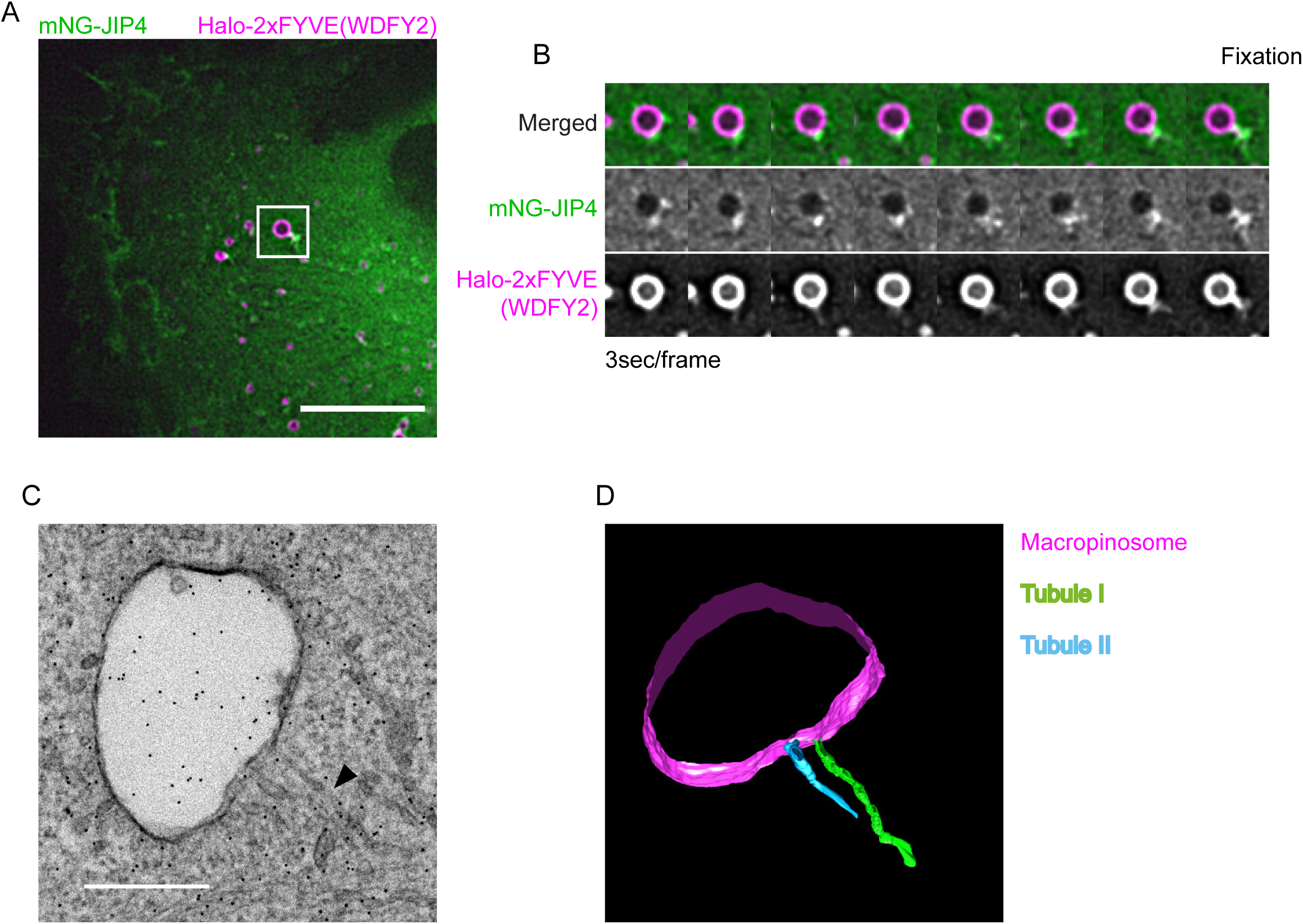
JIP4 tubules are extruded and continuous with macropinosomes. A) Image of RPE1 cell expressing mNeonGreen-JIP4 and Halo-2xFYVE(WDFY2), imaged live during preparation of the CLEM specimen. Scale bar is 5µm. B) Timelapse montage of the tubulating macropinosome until glutaraldehyde fixation. C) Electron micrograph of the macropinosome depicted in (A) and (B). Black dots are gold fiduciaries for electron tomography. The longest tubule emanating from the JIP4 concentration is marked with a black arrowhead. Scale bar is 500nm. D) Model reconstructed from electron tomograph of the macropinosome depicted in (C). The limiting membrane of the macropinosome is in magenta, two separate emanating tubules are in green and blue. The green tubule corresponds to the tubule indicated in (C).

To characterize these tubular structures in detail, we examined the localization of JIP4 together with different markers of membrane tubules. JIP4 tubules emerged from actin-rich domains on the macropinosome (Figure 6A), which were also positive for the actin binding protein Coronin1B (Figure 6B) and the large GTPase Dynamin2 (Figure 6C). JIP4-positive tubules also colocalized with subunits of the retromer complex, VPS26 and VPS35 (Figure 6D, E). The v-SNARE VAMP3, which is sorted into retromer-positive endosomal tubules, also colocalized with JIP4 (Figure 6F). Taken together, this indicates that JIP4 preferentially labels retromer-containing tubules, suggesting that it could be involved in retromer-dependent trafficking.

**Figure 6:**
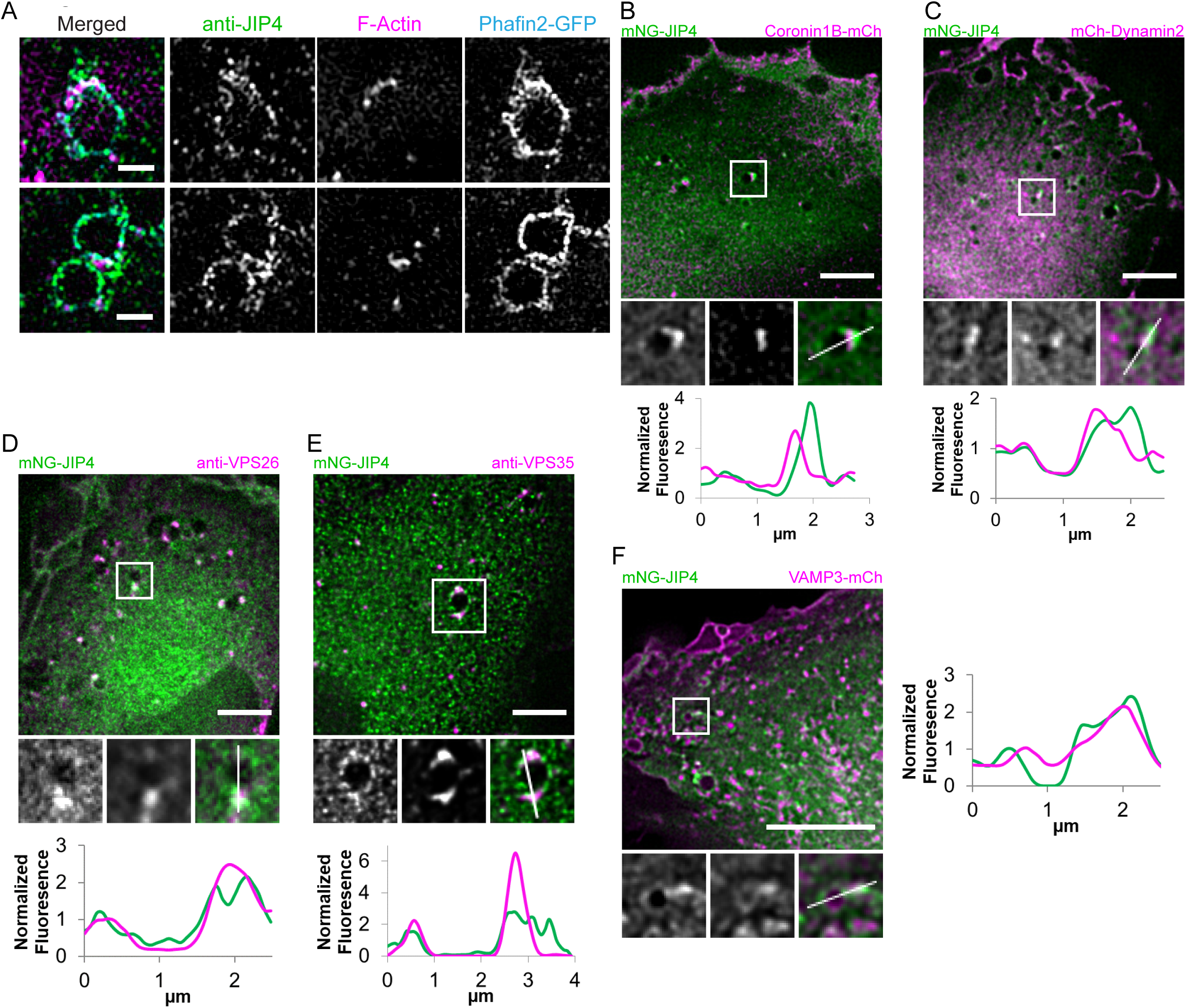
JIP4 tubules bear markers of membrane recycling zones. A) Structured illumination microscopy (SIM) images from two cells expressing Phafin2-GFP, fixed, immunostained against JIP4 and stained for F-actin with phalloidin. Scale bar is 1µm. B) Representative image of RPE1 cell expressing mNeonGreen-JIP4 and Coronin1B-mCherry. Line profile taken along the indicated line from left to right. C) Representative image of RPE1 cell expressing mNeonGreen-JIP4 and Dynamin2-mCherry. Line profile taken along the indicated line from left to right. D) Representative image of RPE1 cell expressing mNeonGreen-JIP4, fixed and immunostained for VPS26. Line profile taken along the indicated line from top to bottom. E) Representative image of RPE1 cell expressing mNeonGreen-JIP4, fixed and immunostained for VPS35. Line profile taken along the indicated line from top to bottom. F) Representative image of RPE1 cell expressing mNeonGreen-JIP4 and VAMP3-mCherry. Line profile taken along the indicated line from left to right. Scale bars in B, C, D, E, and F are 5µm.

In order to investigate the role of JIP4 in trafficking, we next analyzed the phenotype of the JIP4 knockout cells. We measured tubulation from Phafin2-positive macropinosomes (Figure 7A-D) in wild-type and JIP4 knockout cells expressing Phafin2. In addition, cells were transfected with either an empty vector or a JIP4 expressing plasmid. In order to gain a quantitative measurement of tubulation, we measured the co-efficient of variation of the Phafin2 fluorescence over the limiting membrane of the macropinosome (Figure 7C). A higher variation of the fluorescence corresponds to more tubulation events, as these form bright nucleation spots directly at the limiting membrane. (Figure 7B, C). We found that, in comparison to wild-type cells, JIP4 knockout cells showed a small, but significant reduction of macropinosome tubulation in response to Phafin2 expression. In contrast, expression of both Phafin2 and JIP4 in wild-type and knockout cells led to a strong increase in macropinosome tubulation, suggesting that Phafin2 and JIP4 can act together to drive tubulation.

**Figure 7:**
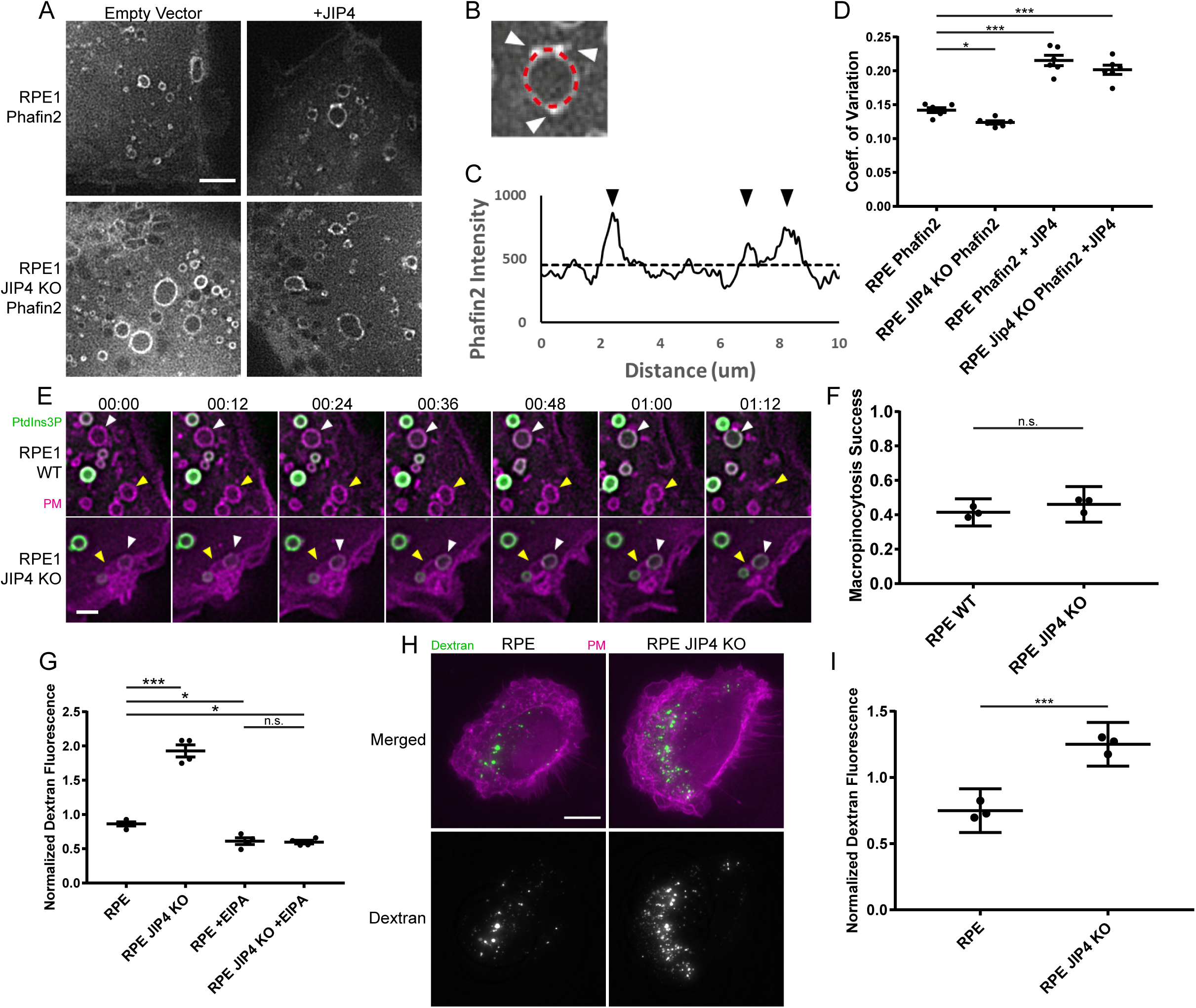
JIP4 promotes tubulation from macropinosomes. A) Representative images of RPE1 cells of the indicated genotypes expressing the specified constructs. The Phafin2 channel is shown. Scale bar is 5µm. B) Example macropinosome, in the Phafin2 channel, depicting the measurement of Phafin2 fluorescence intensity along the limiting membrane of the macropinosome in red dashed line. White arrowheads indicate Phafin2 accumulation at tubule nucleating spots. Note the tubule beginning to extend from the nucleating spot on the top right. C) Line profile of Phafin2 fluorescence intensity taken along the line marked in (B). Black arrowheads correspond to the Phafin2 accumulations in (B). Black dotted line indicates mean of lineplot. D) Coefficient of variation of Phafin2 fluorescence intensity along lineplots taken around macropinosomes >1µm in diameter as in (B), of the indicated genotypes. Mean of 6 experiments shown +/- 95% C.I. (21-72 macropinosomes per condition per experiment) E) Timelapse images of RPE1 cells of the indicated genotypes expressing mNeonGreen-2xFYVE as a PtdIns3P marker and Myrpalm-mCherry as a plasma membrane marker. The yellow arrowheads indicate a macropinosome that fails to mature to an early macropinosome and re-fuses with the plasma membrane. The white arrowheads indicate a macropinosome that matures to an early macropinosome and acquires 2xFYVE. F) Fraction of macropinosomes per cell that successfully mature into an early macropinosome. Mean of 3 experiments shown. Error bars are 95% C.I. (10-15 cells per genotype per experiment) G) Median fluorescence of 20000 cells of the indicated genotype/treatment after 30min uptake of fluorescent 10kDa dextran, measured by flow cytometry. Mean of 4 experiments shown +/- 95% C.I. H) Representative images of RPE1 cells of the indicated genotype after 30min uptake of fluorescent dextran. A plasma membrane marker (fluorescent Wheat Germ Agglutinin) is shown in magenta. I) Total dextran fluorescence per cell of the indicated genotypes after a 30min uptake of fluorescent dextran. Mean of 3 experiments shown +/- 95% C.I. (15-20 cells per genotype per experiment)

We have previously shown that Phafin2 is involved in nascent macropinosome formation [16], and JIP3 and JIP4 have previously been proposed to influence macropinocytosis [17]. We therefore tested if JIP4 is required to form macropinosomes from membrane ruffles. By tracking individual macropinosomes and measuring if they successfully matured into early macropinosomes, we found that loss of JIP4 did not affect early steps of macropinocytosis (Figure 7E, F). This is in line with the localization of JIP4, which only arrives at the macropinosome after maturation into an early macropinosome.

To measure fluid-phase uptake, we performed dextran uptake assays using both flow cytometry and fluorescence microscopy. Using both assays, we noted that JIP4 knockout cells showed significantly elevated intracellular dextran levels in comparison to wild-type cells (Figure 7G, H, I) after a 30min uptake period. We therefore asked if elevated levels of dextran could be detected in different compartments of the endocytic pathway. To this end, we generated stable cell lines expressing RAB5 or LAMP1 in WT and JIP4 KO cells and measured dextran intensity within these compartments. In line with our previous findings, we observed increased dextran fluorescence in both Rab5 (Figure 8A, B) and LAMP1-positive (Figure 8C, D) vesicles, suggesting that more dextran is retained in endolysosomal vesicles in the absence of JIP4. In light of our observation that JIP4 KO cells do not show higher success rates of macropinosome formation and JIP4 does not localize to forming macropinosomes, this elevated intracellular dextran levels could be the result of reduced recycling from macropinosomes. This would be in line with the localization of JIP4 to retromer-containing macropinosome tubules.

**Figure 8:**
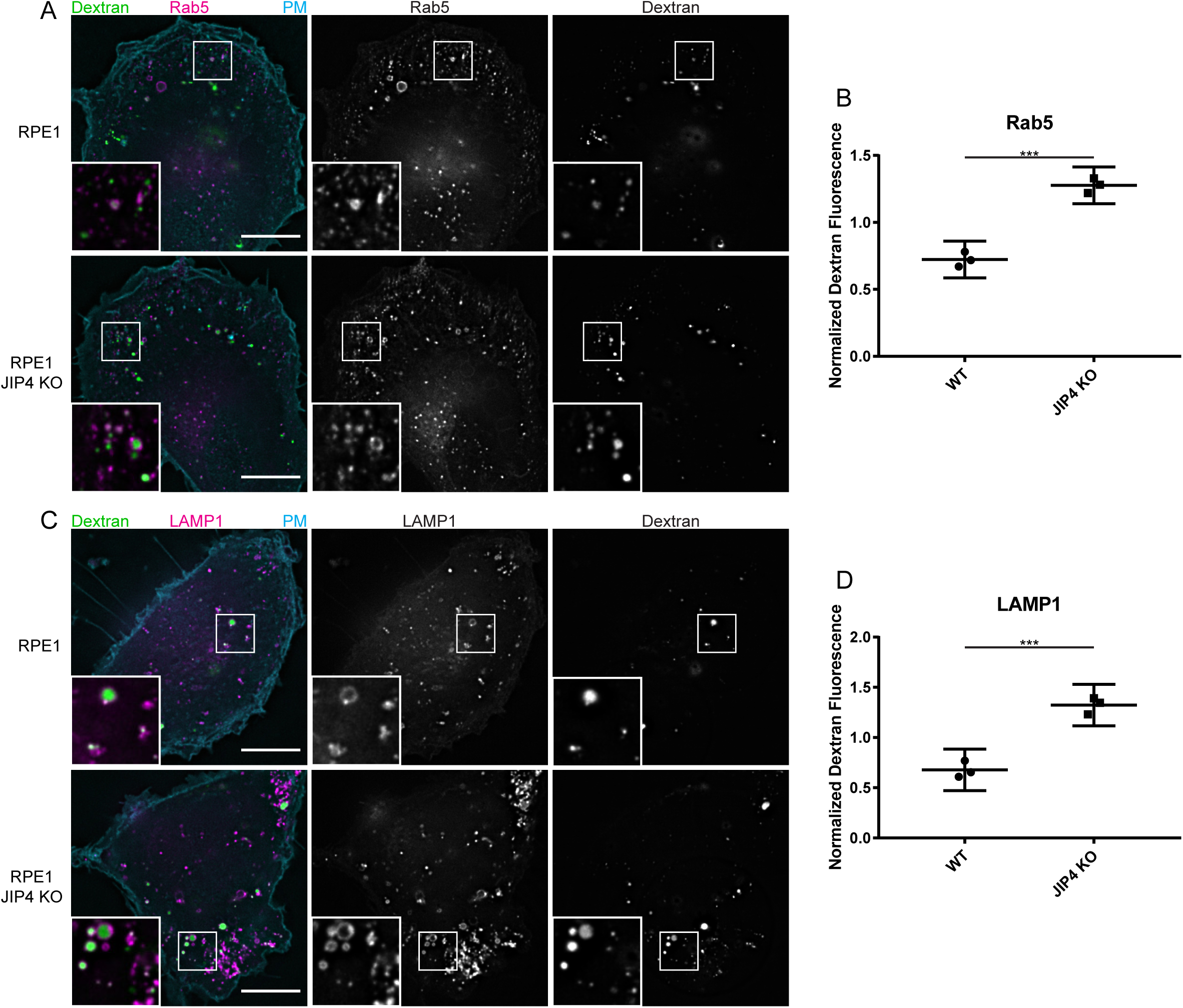
Increased dextran retention in JIP4 KO cells. A) Representative images of RPE1 cells of the indicated genotypes expressing mCherry-Rab5 after a 30min uptake of fluorescent dextran. Scale bar is 5µm. B) Dextran fluorescence in Rab5 positive compartments per cell of the indicated genotype after 30min uptake of fluorescent dextran. Mean of 3 experiments shown +/- 95% C.I. (15-20 cells per genotype per experiment) C) Representative images of RPE1 cells of the indicated genotypes expressing mCherry-LAMP1 after a 30min uptake of fluorescent dextran. Scale bar is 5µm. B) Dextran fluorescence in LAMP1 positive compartments per cell of the indicated genotype after 30min uptake of fluorescent dextran. Mean of 3 experiments shown +/- 95% C.I. (15-20 cells per genotype per experiment)

Taken together, we report a novel, dynamic localization of JIP4, which depends on the lipid-binding protein Phafin2 on macropinosomes. JIP4 localizes to retromer-positive recycling tubules and is required for efficient tubulation.

## Discussion

In the present study, we show that a previously uncharacterized region of JIP4 interacts with the PH domain of the phosphoinositide-binding protein Phafin2, recruiting JIP4 to early macropinosome membranes. Phafin2 binds PtdIns3P generated by the PtdIns 3-kinase VPS34 through its FYVE domain, which localizes it to endosomes and macropinosomes [24, 25]. Our data show that genetic ablation of Phafin2 or the removal of PtdIns3P disrupt the localization of JIP4 to macropinosomes. The recruitment of JIP4 by Phafin2 to membranes does not require other protein or lipid components found on macropinosomes, apart from that needed to anchor Phafin2 to the membrane. The JIP4 homolog JIP3 is not recruited by Phafin2, nor is the Phafin2 homolog Phafin1 capable of recruiting either JIP3 or JIP4. Consistent with this specificity of Phafin2 for JIP4, the ablation of JIP4 did not interfere with the successful completion of macropinocytic internalization, in contrast to JIP3 which was reported to assist macropinosomes in moving through cortical actin [17].

We find that JIP4 is enriched at subdomains of the macropinosome from which membrane tubules are generated and that down- or up-regulating JIP4 levels suppresses or promotes tubulation, respectively. In line with previous studies that functionally implicate JIP4 in endocytic recycling [26], these JIP4 positive tubules contain transmembrane cargo, components of Retromer (a key endocytic recycling complex) [27], and emanate from actin-enriched subdomains on the macropinosome. JIP4 knockout cells retained more of the fluid-phase marker dextran after macropinocytic uptake and this increased cargo retention was found in both early (RAB5) and late macropinosome (LAMP1) compartments.

While Phafin2 shows a biphasic localization to macropinosomes, once to nascent macropinosomes, and another to early macropinosomes [16], JIP4 only binds to Phafin2 at the early macropinosome stage. This suggests that the interaction site between Phafin2 and JIP4 might be inaccessible during the first phase of Phafin2 localization to macropinosomes. Our data do not exclude the possibility that other proteins may contribute to JIP4 localisation, perhaps in a combinatorial manner. Indeed, the binding site for Phafin2 on JIP4 is distinct from those of ARF6 [11], motor proteins, and RAB36 [28]. Furthermore, it has been reported that the JIP3 ARF6-binding-domain only recognises clathrin-coated vesicles after uncoating [26]. While macropinosomes do not use clathrin, newly formed macropinosomes are coated in F-actin [16]. The steric hindrance mechanism proposed by Montagnac et al. for JIP3/ARF6 may therefore also apply to macropinosomes and the JIP4/Phafin2 recruitment.

We additionally observed that this interaction is specific for JIP4 and Phafin2. Phafin2 does not interact with JIP3, nor does Phafin1 bind to JIP4. This is important to note, since several other studies have proposed that JIP3 and JIP4 have overlapping functions, and some phenotypes are reported under double knockdown or knockout conditions [10, 12, 17, 20]. In comparative structural and biochemical analysis, the similarity of the first two coiled-coil regions has been noted [11, 17]. Our data show that the Phafin2 recruitment mechanism distinguishes between the two isoforms. Likewise, despite the high sequence similarity between the Phafin1 and Phafin2 PH domains, only Phafin2 is competent to recruit JIP4.

We find that JIP4 does not localize to the whole macropinosome membrane, but preferably to tubules positive for retromer markers. This is in line with a previous study which described JIP4 localization to late endosomes in close proximity to WASH, which organizes actin on retromer tubules and which reported that JIP3 and JIP4 are required for recycling of the matrix metalloprotease MT1-MMP via endosomal tubules [12].

Based on the described binding of JIP4 to motor proteins, it is tempting to speculate that the tubular localization of JIP4 might couple these membranes to the cytoskeleton and thereby drive tubule formation. Indeed, expression of Phafin2 in JIP4 knockout cells did result in reduced tubulation, whereas expression of both Phafin2 and JIP4 strongly enhanced tubulation. This suggests that Phafin2 and JIP4 act together to enhance tubulation from macropinosomes.

In our previous work, we found that Phafin2 is required during initial steps of macropinosome formation, and that loss of Phafin2 reduces macropinocytosis [16]. While JIP3 and JIP4 have been proposed to play a role in macropinocytosis, we did not observe any defects in macropinocytic fluid-phase uptake in cells deleted for JIP4. In contrast, we did observe enhanced intracellular levels of dextran in JIP4 KO cells, suggesting that these retain more dextran within the cell. This is in line with our observation that Phafin2 is required in early steps of macropinocytosis, whereas JIP4 recruitment only occurs after macropinosomes have successfully entered the cell and have matured into early macropinosomes The increased intracellular dextran levels are consistent with a role of JIP4 in the formation of recycling carriers from macropinosomes.

## Acknowledgements

We thank the Flow Cytometry Core Facility and the Advanced Light Microscopy Core Facility of Oslo University Hospital for technical assistance and access to instruments. We thank Eva Rønning for technical support with yeast two-hybrid assays, Trine Håve for technical support for the APEX2 and the LAPtag experiments, and Ulrikke Dahl Brinch for technical support in generating the low expressing Phafin2-Halo RPE1 cell line. We are grateful to members of the Stenmark Lab for discussions. We are grateful to Philippe Chavrier for sharing the JIP3 plasmid. K.O.S. was supported by a career fellowship from the South-Eastern Norway Regional Health Authority. H.S. was supported by an advanced grant from the European Research Council (project number 788954) and by research grants from the Norwegian Cancer Society and the South-Eastern Norway Regional Health Authority. This work was partly supported by the Research Council of Norway through its Centres of Excellence funding scheme, project number 262652.

## Author contributions

K.O.S and H.S. supervised the study and V.N. co-supervised the study. K.W.T and K.O.S conceived the study and designed experiments. K.W.T generated construct, lentivirus vectors and stable cell lines, performed all live cell imaging, two-hybrid experiments, immunoprecipitation experiment, image analysis and quantifications. K.O.S performed the initial two-hybrid experiments, generated constructs and helped with SIM imaging. V.N. generated stable cell lines, analyzed data and designed experiments. C.C performed APEX2- and LAP-Trap experiments and analyzed mass spectrometry data. A.B. performed electron microscopy. K.W.T, K.O.S. and H.S wrote the manuscript with input from all co-authors.

## Materials and Methods

### Constructs, Cells and Culture Conditions

hTERT-RPE1 cells (ATCC CRL-4000) were grown in DMEM/F12 medium (Gibco) with 10% Fetal Bovine Serum, 5U/ml penicillin and 50µg/ml streptomycin. HeLa cells were grown in DMEM (Gibco) with 10% Fetal Bovine Serum, 5U/ml penicillin and 50µg/ml streptomycin. Cell lines stably expressing constructs were generated by lentiviral transduction at low multiplicity of infection and subsequent antibiotic selection for integration of the expression cassette. The following antibiotics were used: Puromycin (2.5-5µg/ml), Blasticidin (10µg/ml), Geneticin (500µg/ml). VSV-G pseudotyped lentiviral particles were packaged using a third-generation lentivirus system in Lenti-X cells. All lentiviral constructs except Phafin2 were expressed from a phospho-glycerate kinase (PGK) promoter. LAP-tag fusions of Phafin2 were expressed under control of the PGK promoter, whereas other tagged Phafin2 constructs were expressed from an elongation-factor-1α (EF1α) promoter. Transfections were carried out using Fugene 6 (Promega) at a ratio of 3µl reagent per µg DNA. Halotag fusion proteins were labelled with Janelia Fluor 646 Halotag Ligand (Promega) for live cell imaging, or with Janelia Fluor 549 Halotag Ligand (Promega) for correlative light and electron microscopy.

### Generation of JIP4 knockout cell lines

The gRNA sequence (CCTGGACTCGGTGTTCGCGC) was cloned into pX458 with GFP replaced with iRFP. The construct was nucleofected into hTERT-RPE1 cells (Lonza) and sorted by flow cytometry into single cells in a 24 well plate. The resulting colonies were assayed by Western blot and sequencing clones from a genomic PCR flanking the predicted Cas9 cleavage site. The PCR primers for the genomic PCR were CTGGAGGACGGTGTGGTGTA and CGCTCGTACTGGGTGATGAG, with a product length of 266bp, which was cloned into pJet (ThermoFisher Scientific) for Sanger sequencing. Two cell lines lacking JIP4 expression by Western Blot were obtained, and genomic PCR showed one of them to have a G and a C frameshifting insertion. The other clone only produced products with a C frameshifting insertion. The cell line with both alleles containing a confirmed frameshift was chosen for subsequent use, and further validated by immunofluorescence. Sanger sequencing chromatograms, western blot results and immunofluorescence images are shown in Supplemental Figure 1.

### Antibodies

The following antibodies were used.

| Target | Assay | Concentration | Company | Catalog No. |
| --- | --- | --- | --- | --- |
| JIP4 | Western Blot | 1:1000 | Cell Signaling | 5519 |
| Phafin2 | Western Blot | 1:1000 | Sigma-Aldrich | HPA024829 |
| GFP | Western Blot | 1:1000 | Roche | 11814460001 |
| GFP | Immunofluorescence | 1:100 | Roche | 11814460001 |
| JIP4 | Immunofluorescence | 1:100 | Cell Signaling | 5519 |
| EEA1 | Immunofluorescence | 1:160000 | Monash University | Ban-Hock Toh |
| VPS26 | Immunofluorescence | 1:100 | Abcam | ab23892 |
| VPS35 | Immunofluorescence | 1:100 | Abcam | ab10099 |

### Plasmids

JIP4 was obtained by PCR from cDNA reverse transcribed with Superscript IV (Life Technologies) prepared from RPE1 cells. Various constructs of JIP3 were cloned from pEGFP-JIP3, a gift from Philippe Chavrier. VAMP3 was cloned from pEGFP-VAMP3 (Addgene 42310), which was a gift from Thierry Galli [29]. Coronin1B-mCherry (Addgene 27694) and Dynamin2-mCherry (Addgene 27689) were gifts from Christien Merrifield [30]. pX458 (Addgene 48138) was a gift from Feng Zhang [31]. Other constructs were cloned using standard molecular biology techniques.

### Immunoprecipitation

hTERT-RPE1 cells stably expressing GFP or GFP-JIP4 were grown in 6cm dishes up to 80% confluence, washed once with PBS and lysed in lysis buffer (25mM HEPES pH 7.5, 100mM NaCl, 1mM DTT, 0.5% IGEPAL, 1x cOmplete protease inhibitor (Roche), 1x phosphatase inhibitor 2 (Merck) and 1x phosphatase inhibitor 3 (Merck). Cell debris was removed by pelleting at 5000g for 10mins. GFP-Trap beads were added and gently mixed for 2 hours at 4°C. Beads and supernatant were magnetically separated and beads were washed four times with lysis buffer before final denaturation with 1x Laemmli Buffer at 100°C for 20mins.

For tandem affinity purifications, hTERT-RPE1 cells stably expressing LAP or LAP-Phafin2 were grown in 15cm dishes up to 80% confluency. Cells were stimulated with Hepatocyte Growth Factor (HGF) (Merck) at 50ng/ml for 10mins before the experiment. Cells were lysed in lysis buffer (50 mM HEPES pH7.5, 0.1 % NP40, 150 mM KCl, 1 mM EGTA, 1mM MgCl_2_, 1 mM DTT, 15 % Glycerol), cleared by centrifugation at 20,000g for 20 mins, and incubated with GFP-Trap beads for 2 hours. Following 4 washes in lysis buffer, the GFP-Trap bead bound fraction was incubated with recombinant TEV (Merck) overnight at 4°C. The supernatant fraction was collected and incubated with S-protein beads (Merck) for 2 hours. Bound fractions were washed 4 times in lysis buffer and processed for mass spectrometry analysis.

For APEX2 proximity labeling proteomics, hTERT-RPE1 cells stably expressing APEX2-mCitrine-Phafin2 fusions or control fusions were grown in 15cm dishes to 80% confluency. Cells were incubated for 3 hours in 500 µM Biotin-Phenol (Iris) at 37°C, washed in PBS and incubated for 2 min in 2 mM H_2_O_2_ (Merck) at room temperature, and subsequently washed 4 times in Quencher solution (5 mM Trolox (Merck), 10 mM Na-Ascorbate (Merck)). Cells were lysed on ice in RIPA buffer (50mM Tris HCl (pH7.5), 150mM NaCl, 1% Triton X-100, 0.1% SDS, 0.5% NaDOC, 5mM EDTA, 1mM DTT) supplemented with protease inhibitors and 10 mM Na-Ascorbate, cleared by centrifugation at 20,000g for 20 min, and passed through desalting columns to deplete free biotin-phenol. Lysates were subsequently incubated for 2 hours at 4°C with Streptavidin Dynabeads (Invitrogen M-280), and beads were successively washed with RIPA (2 times), PBST (2 times), 1% SDS (2 times), 4 M Urea (2 times), and PBS (5 times) before being processed for mass spectrometry analysis.

### LC–MS/MS, protein identification, and label-free quantitation

Beads containing bound proteins were washed 3 times with PBS, reduced with 10⍰mM DTT for 1⍰h at 56⍰°C followed by alkylation with 30⍰mM iodoacetamide in final volume of 100⍰µl for 1⍰h at room temperature. The samples were digested over night with Sequencing Grade Trypsin (Promega) at 37⍰°C, using 1.8⍰µg trypsin. Reaction was quenched by adding 1% trifluoracetic acid to the mixture. Peptides were cleaned for mass spectrometry by STAGE-TIP method using a C18 resin disk (3M Empore)49. All experiments were performed on a Dionex Ultimate 3000 nano-liquid chromatography (LC) system (Sunnyvale CA, USA) connected to a quadrupole—Orbitrap (QExactive) mass spectrometer (ThermoElectron, Bremen, Germany) equipped with a nanoelectrospray ion source (Proxeon/Thermo). For liquid chromatography separation we used an Acclaim PepMap 100 column (C18, 2⍰µm beads, 100⍰Å, 75⍰μm inner diameter) (Dionex, Sunnyvale CA, USA) capillary of 25⍰cm bed length. The flow rate used was 0.3⍰μL/min, and the solvent gradient was 5% B to 40% B in 120⍰min, then 40–80% B in 20⍰min Solvent A was aqueous 2% acetonitrile in 0.1% formic acid, whereas solvent B was aqueous 90% acetonitrile in 0.1% formic acid.

The mass spectrometer was operated in the data-dependent mode to automatically switch between mass spectrometry (MS) and MS/MS acquisition. Survey full scan MS spectra (from m/z 300 to 1750) were acquired in the Orbitrap with resolution R⍰=⍰70,000 at m/z 200 (after accumulation to a target of 1,000,000 ions in the quadruple). The method used allowed sequential isolation of the most intense multiply charged ions, up to ten, depending on signal intensity, for fragmentation on the higher-energy C-trap dissociation (HCD) cell using high-energy collision dissociation at a target value of 100,000 charges or maximum acquisition time of 100⍰ms. MS/MS scans were collected at 17,500 resolution at the Orbitrap cell. Target ions already selected for MS/MS were dynamically excluded for 45⍰s. General mass spectrometry conditions were: electrospray voltage, 2.0⍰kV; no sheath and auxiliary gas flow, heated capillary temperature of 250⍰°C, heated column at 35⍰°C, normalized HCD collision energy 25%. Ion selection threshold was set to 1e−5 counts. Isolation width of 3.0⍰Da was used.

MS raw files were submitted to MaxQuant software version 1.6.1.0 for protein identification50. Parameters were set as follow: protein N-acetylation, methionine oxidation and pyroglutamate conversion of Glu and Gln as variable modifications. First search error window of 20⍰ppm and mains search error of 6⍰ppm. Trypsin without proline restriction enzyme option was used, with two allowed miscleavages. Minimal unique peptides were set to 1, and false-discovery rate (FDR) allowed was 0.01 (1%) for peptide and protein identification. Label-free quantitation was set with a retention time alignment window of 3⍰min The Uniprot human database was used (downloaded august 2013). Generation of reversed sequences was selected to assign FDR rates.

### Yeast two-hybrid and β-galactosidase assays

Yeast two-hybrid assays were carried out in the yeast strain L40 (ATCC MYA-3332), using LexA and Gal4-Activation Domain (GAD) as paired bait and prey N-terminal fusions [32]. The constructs were co-transformed into yeast and double positive transfectants were selected using leucine + tryptophan drop-out agar medium. Several clones were picked of each condition and pooled to grow overnight liquid cultures for β-galactosidase assay. Liquid β-galactosidase assays were carried out by lysing yeast cells with lysis buffer (100mM Tris HCl pH 7.5, 0.05% Triton-X100) and snap freeze/thaw. Β-galactosidase activity was assayed by hydrolysis of ortho-nitrophenyl-β-galactoside to ortho-nitrophenol in reaction buffer (100mM sodium phosphate buffer pH 7.0, 10mM KCl, 1mM MgSO_4_) at 37°C. The reaction was stopped by addition of a sodium carbonate buffer (250mM final concentration) and immersion in ice as soon as a yellow colour was seen. Ortho-nitrophenol product was quantitated by absorbance at 420nm, reaction rate was calculated, and normalized against quantity of yeast cells (absorbance at 600nm of raw lysate). All experiments were assayed in technical duplicates and 3 separate experiments were carried out for each datapoint reported.

### Immunocytochemistry

hTERT-RPE1 cells of the indicated genotype were grown on glass coverslips. The cells were washed once with ice-cold phosphate buffered saline (PBS) and pre-permeabilized for 5min with PEM buffer (80mM PIPES pH 6.8, 5mM EGTA, 1mM MgCl_2_) containing 0.05% saponin on ice. The cells were then fixed for 20mins on ice with 4% paraformaldehyde in PBS and stained with primary antibody at the listed concentration overnight at 4°C in PBS containing 0.05% saponin. Secondary antibody staining was carried out for 1hr at room temperature in PBS containing 0.05% saponin. Samples were mounted in Mowiol for normal immunofluorescence, and in ProLong Diamond (ThermoFisher Scientific) for Structured Illumination Microscopy.

### Live Cell Microscopy

Live-cell imaging was performed on a Deltavision OMX V4 microscope equipped with three PCO.edge sCMOS cameras, a solid-state light source and a laser-based autofocus. Cells were imaged in Live Cell Imaging buffer (Invitrogen) supplemented with 20⍰mM glucose. Environmental control was provided by a heated stage and an objective heater (20–20 Technologies). Images were deconvolved using softWoRx software and processed in ImageJ/FIJI.

### Structured illumination microscopy

hTERT-RPE1 cells stably expressing Phafin2-GFP were fixed and processed as specified for immunocytochemistry. Phalloidin-Alexa Fluor 647 was included in primary and secondary antibody incubations to stain F-actin, anti-GFP and anti-JIP4 was used to stain Phafin2-GFP and endogenous JIP4. Three-dimensional SIM imaging was performed on Deltavision OMX V4 microscope with an Olympus ×60 NA 1.42 objective and three PCO.edge sCMOS cameras and 488nm, 568nm and 647nm laser lines. Cells were illuminated with a grid pattern and for each image plane, 15 raw images (5 phases and 3 rotations) were acquired. Super-resolution images were reconstructed from the raw image files aligned and projected using Softworx software (Applied Precision, GE Healthcare). Images were processed in ImageJ/Fiji.

### Quantifying endogenous JIP4 on Early Macropinosomes

Cells of the listed genotype were processed and fixed for immunocytochemistry. 15 fields of view of each condition were acquired (typically 1-3 cells per field of view) without changing acquisition parameters. EEA1 positive structures of at least 5 pixels were segmented from each image and the mean pixel intensity of each structure in the JIP4 channel was obtained. Each dataset was normalized by the mean of the entire experiment to control for staining and acquisition variation.

### Quantifying JIP4 association to tubules

Cells of the listed genotype stably expressing the 2xFYVE^WDFY2^ probe and mNeonGreen-JIP4 were stimulated with 50ng/ml HGF to trigger macropinocytosis, imaged live and videos were taken for 5mins at intervals of 3secs. Tubules (membrane deformations that exceeded 6 pixels in length, 80nm/pixel) that formed during that time period were marked in the 2xFYVE^WDFY2^ channel. The cytoplasmic background fluorescence for JIP4 of each cell was estimated by taking a 100×100 pixel square and measuring the mean fluorescence in the JIP4 channel. Each identified tubule was classified as JIP4 positive if it contained JIP4 fluorescence at least 50% over the background fluorescence determined above. Each cell was treated as a single biological datapoint (proportion of tubules JIP4 positive).

### Quantification of Co-efficient of Variation

RPE1 or RPE1 JIP4 KO cells expressing Phafin2-mTurquoise were transfected one day before the experiment with either empty vector or mNeonGreen-JIP4. Cells were stimulated with HGF (50ng/ml) and timelapse images were captured. The image frame corresponding to 30 secs after the start of imaging was extracted and used for further analysis. All macropinosomes greater than 1µm in diameter were included in the analysis. A 3-pixel wide line was manually drawn in ImageJ around each macropinosome such that the entire circumference of the macropinosome was included. ImageJ reports the average gray value of the 3-pixel thickness at each position along the line. These values were used to compute the co-efficient of variation of Phafin2 intensity along the circumference of each macropinosome.

### Measurement of protein fluorescence intensities at the macropinosome membrane

Live cell imaging was performed as described earlier on RPE1 cells expressing the specified proteins. HGF (50ng/ml) was used to trigger macropinocytosis and timelapse videos were captured. Newly formed macropinosomes were identified in timelapse movies and manually tracked by using Phafin2 or membrane markers as reference. For each time point, a region of their limiting membrane was marked as region of interest. Fluorescence intensity of a circular ROI (10 pixel diameter) surrounding the marked region was quantified in all image channels and measurements were exported for further analysis.

### Flow Cytometry – Dextran Uptake

Cells were seeded in 6 well plates at a density of 1×105 the day before the experiment. The media was replaced by prewarmed media containing 0.5mg/ml dextran-Alexa Fluor 488 (10kDa) and 50ng/ml HGF (and EIPA where indicated) and cells were incubated at 37°C for 30mins. After the incubation, cells were washed five times with prewarmed media, trypsinized, and placed on ice after neutralization of trypsin. Flow cytometry was performed shortly after trypsinization with an LSRII flow cytometer (BD Biosciences).

### Dextran Fluorescence by Microscopy

Cells of the indicated genotypes were seeded in glass-bottomed mattek dishes. The media was replaced by prewarmed media containing 0.5mg/ml dextran-Alexa Fluor 488 (10kDa) and 50ng/ml HGF. Cells were incubated at 37°C for 30mins. After the incubation, cells were quickly washed four times with prewarmed media, once with phosphate buffered saline, and fixed for 10min at room temperature using 4% paraformaldehyde in PBS. The cells were gently washed three times in PBS and the plasma membrane labelled with Wheat Germ Agglutinin-Alexa Fluor 647 (Molecular Probes) at 5µg/ml for 10mins in PBS. The cells were washed twice, the nuclei labeled with Hoescht 33342 (Molecular Probes), and imaged in PBS. Image z-stacks of 6µm were acquired at an interval of 250nm and deconvolved. One cell was measured per field of view acquired (the field of view was typically only large enough to fully fit one cell). For whole cell dextran fluorescence measurements, image stacks were z-projected using the sum of intensities. Cell outlines were manually traced in ImageJ using the plasma membrane marker as a guide. Background values (compensation for residual nonspecific dextran and imperfect deconvolution) were obtained from a 100×100 pixel square outside cells and subtracted from the fluorescence measured inside the cells. For organelle specific values, the image plane that was most in focus was extracted from the stack. Organelles of at least 5 pixels (approximately diffraction limit of 240nm) were segmented using the listed organelle marker and the fluorescence measured. Values reported are computed per cell. Each experiment was normalized by the average of all datapoints in that experiment to account for acquisition parameters (these were held constant for all image stacks acquired in an experiment).

### Correlative Light and Electron Microscopy

Cells were seeded on gridded Matteks the day before the experiment. Light microscopy was carried out as specified in “Live Cell Microscopy” with timelapse acquisition while cells were stimulated with 50 ng/ml HGF. Directly after live cell imaging fixation was carried out using a final concentration of 2% glutaraldehyde in 0.1 M PHEM buffer (80 mM PIPES, 25 mM HEPES, 2 mM MgCl_2_, 10 mM EGTA, pH 6.9) for 1 h and postfixation was done in 1% OsO_4_ and 1.5% KFeCN in the same buffer (1 h). Samples were further en bloc stained with 4% aquaeus uranyl acetate for 1 h, dehydrated in graded ethanol series and embedded with Epon-filled BEEM capsules (EMS; Polysciences, Inc., 00224) placed on top of the Mattek dish. After polymerization blocks were trimmed down to the regions previously identified on the OMX microscope and now imprinted on the Epon block. 200 nm sections were cut on an Ultracut UCT ultramicrotome (Leica, Germany) and collected on formvar coated slot grids. Samples were imaged using a Thermo ScientificTM TalosTM F200C microscope equipped with a Ceta 16M camera. Single-axes tilt-series for tomography were acquire between −60° and 60° tilt angles with 2° increment. Tomograms were computed in IMOD using weighted back projection [Kremer et al., 1996, PMID:8742726]. 3D modeling was performed by manual tracing of the macropinosome membrane in IMOD software version 4.9.3. Display of tomogram slices was also performed using IMOD software.

### Rapamycin Recruitment

The mitochondrial anchor was constructed by fusing tandem FKBP12 FK506 binding domains to an N-terminal Tom70-derived mitochondrial targeting signal, with mTagBFP2 as localization marker. The FKBP-Rapamycin-Binding (FRB) domain of mTOR with a T2098L stabilization mutation and mNeonGreen was appended to Phafin2 at the C-terminus of Phafin2. The mCherry tagged JIP4 was not further modified. These three constructs were transfected into RPE1 cells as previously described and images acquired in live timelapse microscopy. A final working concentration of 10µM of SAR-405 was used to dissociate Phafin2 from vesicles, and a final working concentration of 250nM of rapamycin was used to recruit tagged Phafin2 to the mitochondrial anchor, added 5mins after treatment with SAR-405. Images were acquired before treatment, 5mins after treatment with SAR-405 and approximately 30mins after treatment with rapamycin. Intensity measurements were obtained by segmenting images using the mTagBFP2 mitochondrial marker.

### Statistical Analysis

Statistical analysis was carried out in Graphpad Prism (Graphpad Software). Student’s t-test was used to compare two groups. ANOVA was used to compare multiple groups and Holm-Sidak was used to correct for multiple comparisons. The threshold for significance was set at p=0.05. All comparisons made are reported regardless of significance. In all figures, * indicates that p<0.05, ** indicates that p<0.01, and *** indicates that p<0.001.

## Figure Legends

**Supplemental Figure 1:**
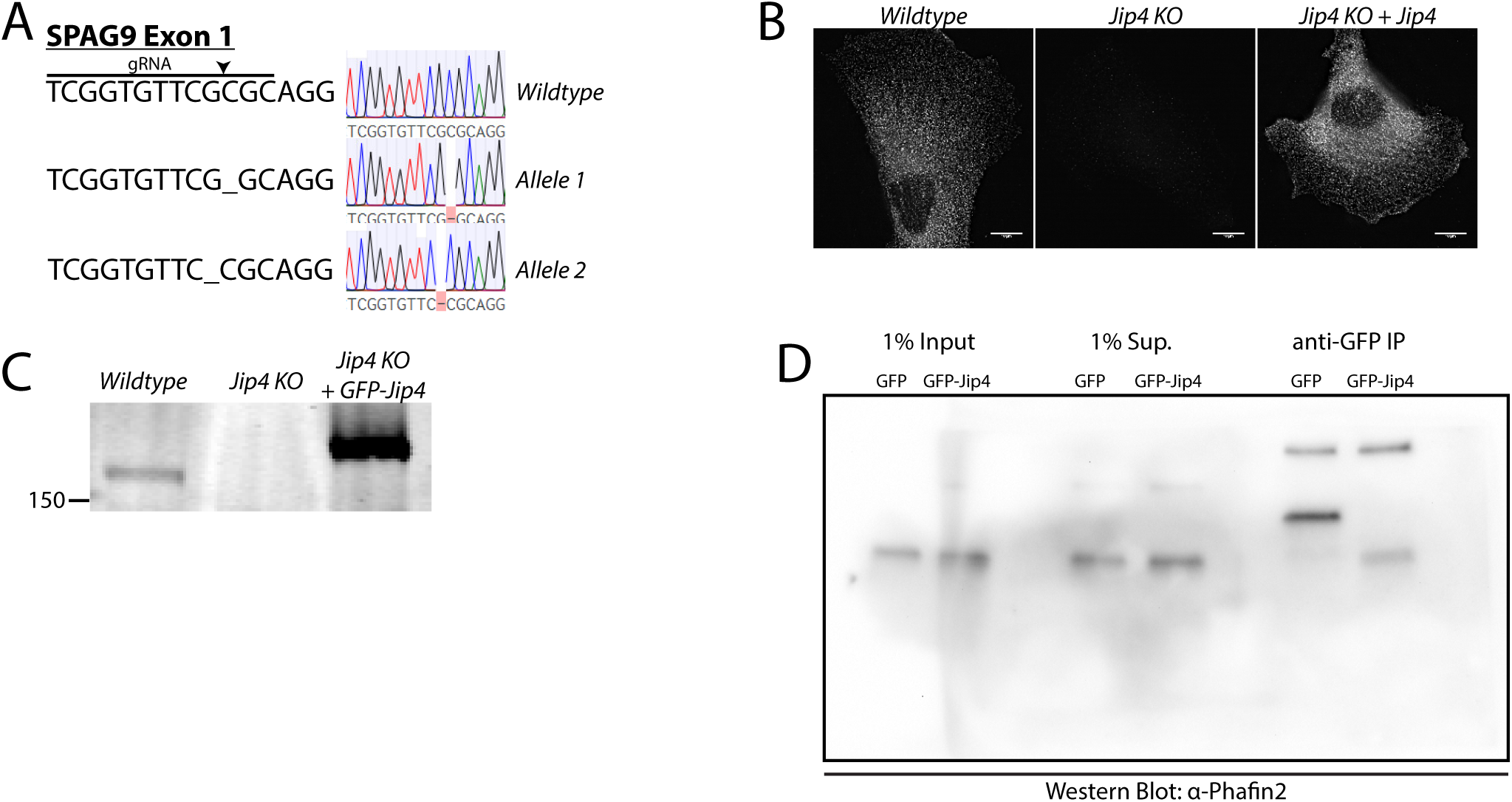
Generation and verification of RPE1 JIP4 knockout cell line. A) Guide RNA for CRISPR/Cas9 knockout. The predicted cut site is indicated. Sanger sequencing chromatograms show different frameshift insertions for both alleles. No wildtype sequencing results were recovered from the JIP4 KO cell line. B) Immunofluorescence using anti-JIP4. Images were acquired at the same settings and presented with equal brightness scaling. C) Western blot using anti-JIP4 on cell lysate from wildtype, JIP4 KO, and JIP4 KO expressing GFP-JIP4.

**Supplementary Video 1: JIP4 localizes to Phafin2 positive early macropinosomes.** Shown is a macropinocytosing RPE1 cell with Phafin2-mTurquoise2 (pseudocolored green) and mNeonGreen-JIP4 (pseudocolored magenta). Note that the nascent macropinosomes entering on the right display a burst of Phafin2 that is not accompanied by JIP4, while the early macropinosomes acquire both Phafin2 and JIP4.

**Supplementary Video 2: JIP4 localizes in subdomains on Rab5 positive macropinosomes.** Shown is a macropinocytosing RPE1 cell with mNeonGreen-JIP4 (pseudocolored green) and mCherry-Rab5 (pseudocolored magenta). JIP4 localizes to dynamic subdomains as the macropinosome acquires Rab5.

